# Localized actin cortex perturbation generates cell-scale membrane tension gradients

**DOI:** 10.64898/2026.05.06.721853

**Authors:** Yogishree Arabinda Panda, Elisabeth Fischer-Friedrich

## Abstract

Membrane tension is a key mechanical regulator of cell signaling, morphology, and division. Whether cells can sustain spatial gradients in membrane tension, or whether such asymmetries are rapidly dissipated by long-range tension propagation, remains actively debated. Here, we use the tension-sensitive fluorescent probe FliptR to directly measure in-plane membrane tension before and after localized optogenetic activation of RhoA in mitotic HeLa cells. We find that localized RhoA activation generates a sustained, cell-scale membrane tension gradient, with tension elevated on the non-activated side. This gradient depends on an intact actin and microtubule cytoskeleton and is accompanied by polarized cytoskeletal remodeling: cortical f-actin enrichment at the activation site and asymmetric microtubule growth on the opposite side. Tether-force measurements reveal enhanced membrane–cortex adhesion at the activated side, with no corresponding increase in in-plane tension, reconciling an apparent discrepancy between prior studies. A coarse-grained membrane chemical potential accounts for gradient maintenance through spatially heterogeneous membrane–cortex coupling. Together, our findings demonstrate that cells can actively generate and sustain spatially patterned mechanical states through localized cytoskeletal signaling.

## I. INTRODUCTION

Plasma membrane tension is a fundamental biomechanical parameter that regulates a wide range of cellular processes, including endocytosis, exocytosis, cell migration, division, and mechanosensing (1–4). By regulating mechanosensitive ion channels and other tension-sensitive membrane proteins, membrane tension can couple local mechanical perturbations to intracellular signaling and has therefore been proposed to act as a cell-wide integrator of mechanical cues (3, 5). Despite its recognized biological importance, the spatial organization of membrane tension within individual cells — and in particular the extent to which tension gradients can be established and maintained — remains poorly understood.

A central question in cell mechanics is whether membrane tension equilibrates rapidly across the cell or whether it can be spatially non-uniform over physiologically relevant timescales. Early theoretical frameworks predicted that, owing to the low viscosity of the lipid bilayer, tension perturbations should propagate rapidly across the plasma membrane (4, 6). However, experimental measurements using optical tweezers and membrane tether pulling revealed that local tension perturbations do not propagate detectably across the cell on timescales of hundreds of seconds, suggesting that friction between the membrane and the underlying cytoskeleton strongly attenuates long-range tension transmission (3, 7). This frictional coupling, arising from membrane-protein obstacles and membrane–cortex attachments, has been incorporated into theoretical models of membrane mechanics (7, 8).

Seemingly at odds with these findings, de Belly et al. (9) showed that localized actin-based protrusions and actomyosin contractions triggers rapid, cell-wide changes in membrane tether force within seconds. This was interpreted as evidence for fast, cortex-mediated propagation of membrane tension across the cell, driven by active cytoskeletal forces rather than passive lipid flow. Together, these observations point to an apparent discrepancy: local membrane-tension perturbations generated directly at the plasma membrane remain spatially confined, whereas cortex-mediated changes in membrane tension appear to equilibrate rapidly across the cell.

To resolve this apparent discrepancy, we argue that tether forces report an apparent membrane tension, *γ*_app_ = *γ*_mem_ + *γ*_adh_, which conflates in-plane lipid tension *γ*_mem_ with membrane–cortex adhesion energy *γ*_adh_ (4, 10). Thus, while de Belly et al. demonstrated cell-wide changes in tether force following localized cortical activation, their measurements did not decouple these two contributions. Whether localized cortical signaling can generate and sustain spatial gradients specifically in in-plane membrane tension, and which cytoskeletal mechanisms underlie such asymmetry, has therefore not been directly addressed.

Resolving this measurement ambiguity requires independent measurements of in-plane membrane tension that are not confounded by membrane–cortex adhesion. The fluorescent membrane tension probe FliptR offers such an approach: its fluorescence lifetime reports changes in lipid packing and membrane order that correlate with in-plane membrane tension (11, 12). This makes FliptR a complementary and orthogonal tool to tether-based methods for probing membrane tension in living cells.

A second unresolved question concerns the role of the cytoskeleton in shaping the spatial distribution of membrane tension. Prior work has established that the actin cortex and myosin-II-driven contractility are major determinants of cortical tension and cell shape (12–15). In mitotic cells, the cortex is particularly well-organized, adopting a nearly isotropic, tension-bearing structure that contributes to cell rounding and the mechanical integrity required for accurate chromosome segregation (12, 15, 16). Yet the specific cytoskeletal mechanisms by which localized cortical signaling could generate and sustain spatial asymmetries in membrane tension across such an initially uniform cortex remain to be identified.

Here, we address these questions by combining localized optogenetic activation of RhoA (17) with FliptR-based membrane-tension readouts, tether-force measurements, pharmacological and live-imaging perturbations of the cytoskeleton in mitotic HeLa cells. We demonstrate that localized RhoA activation generates a sustained, cell-scale gradient in FliptR lifetimes suggesting a corresponding gradient in in-plane membrane tension, with tension elevated on the non-activated side. We show that this gradient is maintained by polarized membrane–cortex adhesion – driven by local actin polymerization at the activation site – and is shaped by asymmetric cytoskeletal remodeling involving both actin and microtubules. Together, our results reconcile the apparent discrepancy between tether-force and FliptR-based measurements suggesting that the cell-wide tether-force changes reported by de Belly et al. (9) primarily reflect spatially patterned changes in membrane–cortex adhesion rather than uniform propagation of in-plane tension. More broadly, these findings support a framework in which spatial variations in membrane–cortex coupling maintain opposing membrane tension gradients, within the framework of an effective membrane chemical potential. Furthermore, this work identifies cytoskeletal polarization as a key driver of membrane tension asymmetries.

## II. RESULTS

### A. Localized optogenetic activation of opto-RhoA induces a sustained plasma membrane tension gradient across mitotic cells

To investigate the membrane tension propagation and the emergence of tension gradients, we employed the fluorescent probe FliptR to quantify spatial variations in membrane tension following localized opto-RhoA (17) activation in transgenic HeLa cells expressing opto-RhoA–mCherry, see Fig. 1a and Materials and Methods. For convenience, we restricted our study to spherical weakly-adherent cells pharmacologically arrested in mitosis, see Materials and Methods, in order to obtain well-controlled spherical cell shapes and an approximately uniform cell cortex (12, 16). Upon blue light exposure, the opto-RhoA construct is recruited to the plasma membrane through the BcLOV4 domain, which acquires membrane affinity under illumination (17) (Fig. 1b).

**Figure 1.**
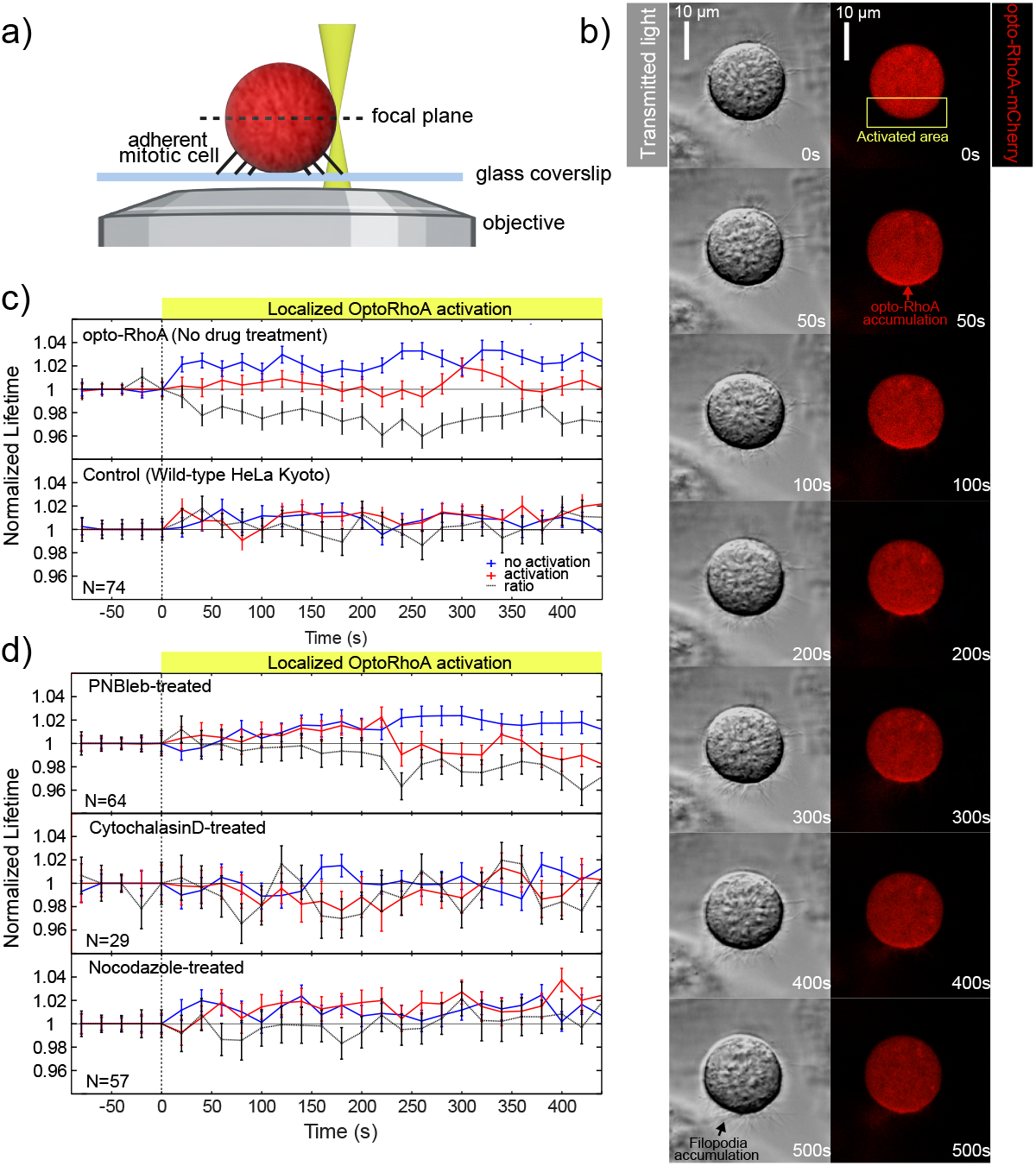
Localized optogenetic RhoA activation induces a sustained membrane tension gradient mediated by actin and microtubules. a) Schematic of the experimental setup for localized opto-RhoA-mCherry activation in mitotically arrested HeLa cells. Images were acquired in the equatorial plane. Cells were arrested in mitosis through incubation with 4 *µ*M STC. b) Time series showing polarized accumulation of cortical opto-RhoA-mCherry upon local blue-light illumination (473 nm). Cortical polarization is further reflected by a localized accumulation of filopodia at the activation site indicated by the black arrow in the left lowermost panel. c,d) Time evolution of normalized FliptR lifetimes in the activated and non-activated half of the cells for different conditions. c) Top panel: Optogenetic activation keeps FliptR lifetimes approximately constant in the activated half of the cell (red curve), while increasing lifetimes in the non-activated half (blue curve). As a result, the lifetime ratio (activated/non-activated, black curve) decreases, suggesting that membrane tension increases and remains persistently elevated in the non-activated half of the cell upon localized opto-RhoA activation. Bottom: In wild-type HeLa cells lacking the optogenetic construct, FliptR lifetimes increase slightly in both cell halves, without the emergence of a tension gradient across the cell. d) Cytoskeletal perturbations delay or abolish the formation of the membrane tension gradient. Top: Myosin inhibition with PN-Blebbistatin (20 *µ*M) delays the onset of the membrane tension gradient. Middle and bottom: Actin depolymerization with Cytochalasin D (4 *µ*M) or microtubule depolymerization with Nocodazole (10 *µ*M) abolishes membrane tension gradient formation. c,d) Data points represent medians across multiple measured cells, and error bars indicate the standard error of the median.

Previous studies have reported fast long-range propagation of apparent membrane tension upon optogenetic activation of RhoA in non-adherent interphase cells (9). In our study, we find surprisingly the formation of a persistent membrane tension gradient across the equatorial plane of the cell using FliptR as a reporter of in-plane membrane tension. Specifically, periodic localized activation of opto-RhoA resulted in a sustained increase in FliptR lifetime at the plasma membrane on the non-activated side, indicative of elevated in-plane membrane tension at the non-activated side, while no significant change was observed in the FliptR lifetime from the plasma membrane on the activated side (Fig. 1c, blue and red curves, top panel). Correspondingly, the ratio of FliptR lifetimes (activated/non-activated) was significantly decreased upon activation (Fig. 1c, black curve, top panel). Concurrently, we observed an asymmetric growth of filopodia-like membrane reservoirs at the activated side, suggesting localized cortex and membrane remodeling, see Fig. 1b, bottom panel.

In contrast, control experiments performed in non-transfected HeLa Kyoto cells subjected to the same illumination protocol showed a modest insignificant increase in FliptR lifetime on both sides, with no significant difference between activated and non-activated regions (Fig. 1c, lower panel). This absence of spatial asymmetry in wild-type cells verifies that the observed tension gradient is specifically driven by opto-RhoA activation rather than light exposure in itself. Together, these results demonstrate that localized optogenetic activation of RhoA is leading to a sustained membrane tension gradient across mitotic cells with tension elevation on the side opposite to activation.

### B. Generation of membrane tension gradients depends on the actin and microtubule cytoskeleton

To assess the contribution of cytoskeletal components to membrane tension propagation and gradient formation, we examined the effect of targeted pharmacological perturbations in opto-RhoA–expressing HeLa cells. Cells were treated with PN-Blebbistatin (myosin II inhibitor), Cytochalasin D (actin polymerization inhibitor), or Nocodazole (microtubule depolymerizing agent) prior to localized opto-RhoA activation. Inhibition of myosin II activity by PN-Blebbistatin attenuated the response to localized opto-RhoA activation, resulting in a delayed and reduced membrane tension gradient (Fig. 1d, top panel). In contrast, disruption of actin polymerization with Cytochalasin D or microtubule integrity with Nocodazole prevented the formation of a detectable membrane tension gradient under identical activation conditions (Fig. 1d, middle and lower panel). These results indicate that membrane tension gradient formation in mitotic cells requires an intact and dynamic cytoskeletal network. While actomyosin contractility contributes to the efficiency and kinetics of gradient establishment, both actin filaments and microtubules are essential for the emergence of sustained spatial asymmetry in membrane tension.

### C. Optogenetic activation of RhoA leads to localized increase of cortical actin as well as enhanced and polarized microtubule growth

To further investigate how actin and microtubule dynamics contribute to membrane tension gradient formation, we performed time-resolved live-cell imaging of cytoskeletal organization using SiR-actin and Tubulin Tracker™ Deep Red during optogenetic activation (see Materials and Methods). Localized activation of opto-RhoA resulted in a pronounced and asymmetric increase in cortical f-actin intensity on the activated half of the cell, accompanied by a relative decrease on the non-activated side (Fig. 2a-c). This redistribution is consistent with localized RhoA-driven actin polymerization at the side of activation.

**Figure 2.**
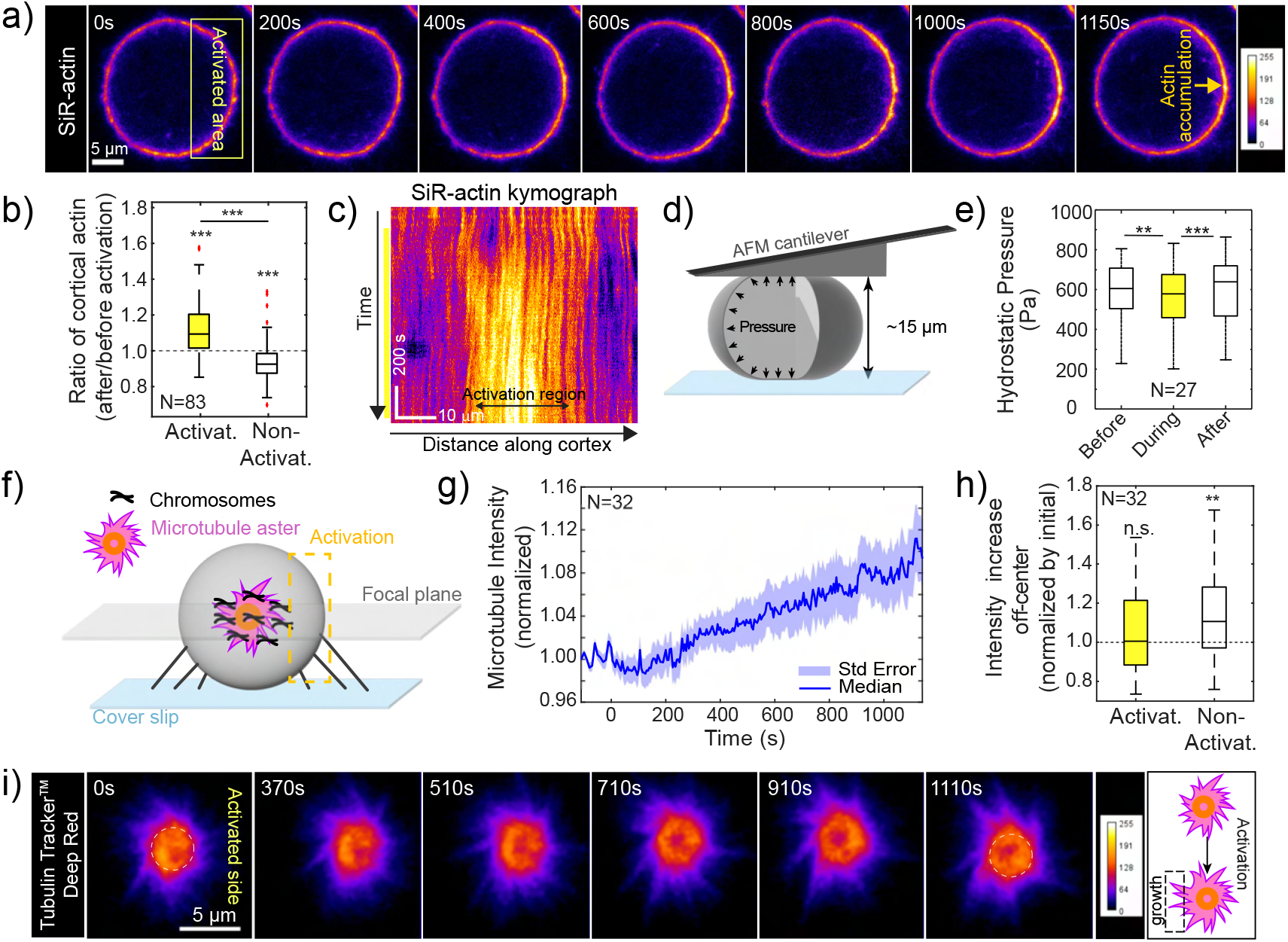
Localized optogenetic activation of RhoA induces f-actin accumulation at the activated cortex and enhanced microtubule growth at the opposite side. a) Time series of cortical f-actin fluorescence (SiR-actin) in a mitotic cell (acquired in the equatorial plane), shows accumulation at the activated side and a depletion at the non-activated side upon sustained optogenetic activation. b) Boxplots showing the ratio of f-actin intensity in the final frame (*t* ≈ 1100 s) relative to the initial frame at both the activated and non-activated cortical sides. Statistical significance between the two distributions was assessed using a paired t-test (*p* ≤ 0.05). In addition, each distribution was tested against a mean of one using a one-sample t-test (*p* ≤0.05). c) Kymograph of SiR-actin intensity showing f-actin accumulation in the activated cortical region (cell as in panel a). d) Schematic of the parallel plate confinement assay setup used to determine intracellular hydrostatic pressure with a wedged AFM cantilever. e) Boxplot showing intracellular hydrostatic pressure before, during, and after localized optogenetic activation in the equatorial plane. f–h) Localized RhoA activation enhances microtubule polymerization and induces asymmetric aster growth. f) Schematic of microtubule imaging in HeLa cells in mitotic arrest using Tubulin Tracker™ Deep Red. The bipolar spindle reorganizes into a characteristic monopolar aster upon mitotic arrest induced by Eg5 inhibition with STC (19). g) Normalized microtubule fluorescence intensity over time, showing increased microtubule polymerization following local RhoA activation. The curve represents the median across all measured cells. Intensities were normalized to the fluorescence signal before activation (negative times). h) Boxplot showing the ratio of microtubule intensity in the final frame (*t* ≈ 1100 s) relative to the initial frame for regions adjacent to the microtubule aster on the RhoA-activated side (left) and the non-activated side (right), see Fig. S1c. i) Time-lapse images showing the evolution of the microtubule aster. The white dashed circles have the same radius in the initial and final images and serve as a reference for the increase in aster size.

Further, we examined whether localized opto-RhoA activation induced directed cortical actin flows toward the stimulated region. Using live imaging of f-actin dynamics, we did not detect any consistent or measurable directed cortical flow of f-actin towards the site of optogenetic activation under our experimental conditions (Fig. 2c). This indicates that the observed increase in f-actin signal in the activated cortex region is not driven by large-scale advective transport of pre-existing actin filaments. Instead, it is more consistent with local actin polymerization and cortex remodeling due to localized enhancement of RhoA-mediated signaling.

To assess whether localized opto-RhoA activation induces an increase of cortical tension and hydrostatic pressure, we measured confinement forces of cells within an AFM-based parallel plate assay (14) (Fig. 2d). Measurement was performed for each cell before, during and after localized opto-RhoA activation revealing a shallow decrease in intracellular hydrostatic pressure upon localized opto-RhoA activation (Fig. 2e), hinting at a reduction of cortical tension on average across the cell. This reduction of pressure was reversible, eventually returning to baseline levels after the localized activation was stopped (Fig. 2e) indicating that the effect is specifically coupled to transient changes in RhoA activity. We speculate that cortical tension and pressure slightly decreased upon RhoA activation (rather than increasing) due to the predicted non-monotonic dependence of cortical contractility on actin cortex thickness and actin filament lengths in an already strongly activated mitotic actin cortex in a mitotic cell (18).

Investigating also microtubule dynamics upon optogenetic activation, we observed an overall increase in microtubule signal in the microtubule aster following activation (Fig. 2f,g), with a significant asymmetric increase on the nonactivated side, while the activated side showed no change in microtubule intensity (Fig. 2h,i). This asymmetry indicates a spatial bias in microtubule reaction kinetics changes across the cell.

In summary, we find polarized redistribution of both the actin and the microtubule cytoskeleton, with f-actin enrichment at the activated side of the cortex and increased microtubule polymerization at the opposite, non-activated cell half. This polarity suggests a coordinated, spatially organized response to localized RhoA signaling.

### D. Membrane-cortex attachment increases at the site of RhoA activation

To better understand how the emerging membrane-tension gradients induced by localized optogenetic activation of RhoA are maintained, we asked whether this process is accompanied by changes in membrane–cortex adhesion. Previous work has shown that membrane tether forces, *F*_th_, depend on membrane–cortex adhesion according to

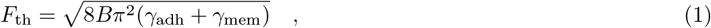

where *γ*_adh_ denotes the membrane–cortex adhesion energy per unit area, *γ*_mem_ the in-plane membrane tension, and *B* the membrane bending stiffness (10). Since membrane tension was not significantly altered on the activated side of the cell according to FliptR measurements (Fig. 1, topmost panel), we reasoned that changes in tether force would primarily reflect changes in membrane–cortex adhesion following localized optogenetic activation.

Using AFM-based tether force measurements in optogenetically activatable cells, we found that localized opto-RhoA activation led to a significant increase in tether force, indicating enhanced membrane–cortex attachment at the site of activation (Fig. 3a,b).

**Figure 3.**
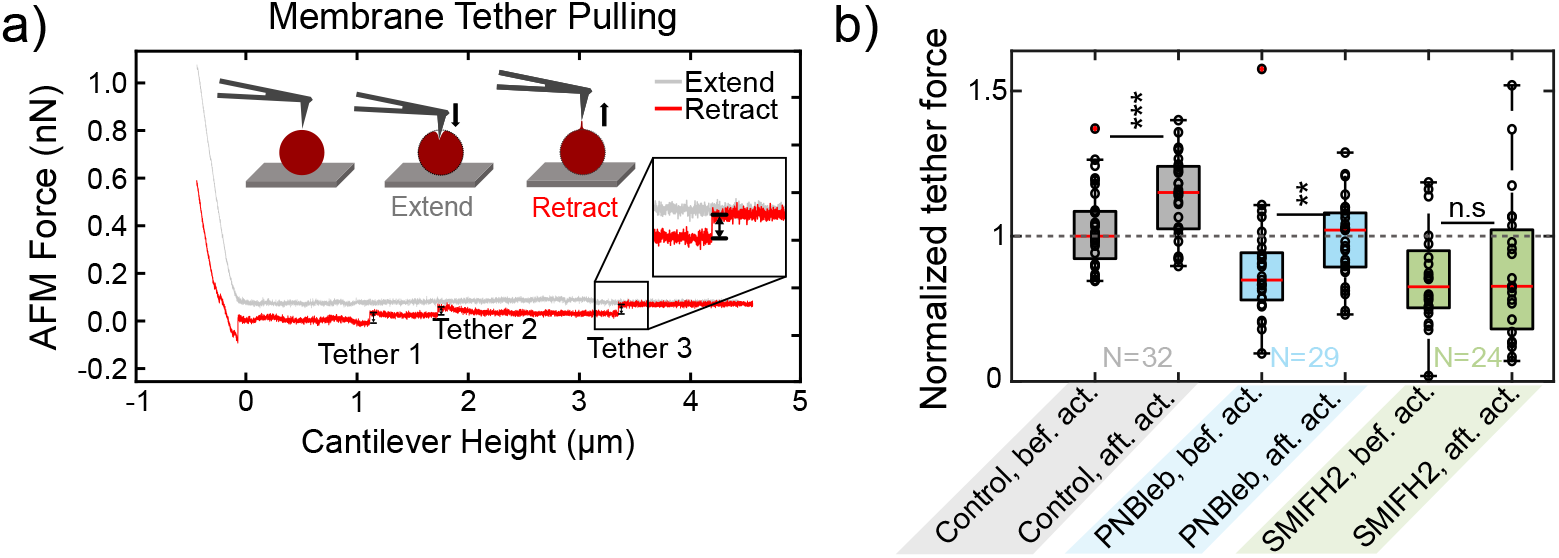
Membrane tether forces increase upon RhoA activation in dependence of formin-mediated actin polymerization but not myosin contractility. a) Exemplary force-indentation curve on a HeLa cell in mitotic arrest showing force steps associated to the membrane tether detachment events. Force step magnitudes are interpreted as tether forces. b) Boxplots showing normalized tether forces measured on HeLa cells expressing the opto-RhoA construct before and after optogenetic activation at the topmost plane of the cell. Cells were measured in control conditions (no drug), treatment with the myosin inhibitor PN-Blebbistatin (blue, 20 *µ*M) and treatment with the formin inhibitor SMIFH2 (green, 40 *µ*M). Each data point represents the median of force steps measured on one cell in the respective condition. Normalization was achieved by dividing all forces through the median force value of the control condition (before activation) of the respective measurement day. Statistical significance between the two distributions was assessed using a paired t-test (*p ≤* 0.05).

To exclude the possibility that this increase in tether force resulted indirectly from enhanced myosin-dependent contractility and elevated hydrostatic pressure, we inhibited myosin II activity using PN-Blebbistatin. Myosin II inhibition did not abolish the activation-induced increase in tether force, suggesting that optogenetically-induced changes of actomyosin contractility are not the main driver of the observed tether-force changes, see Fig. 3b (blue boxplots). In contrast, inhibition of actin polymerization with the formin inhibitor SMIFH2 abolished the increase in tether force upon localized activation, indicating that dynamic actin assembly is required for strengthening membrane– cortex coupling, see Fig. 3b (green boxplots). Together, these results suggest that localized RhoA activation enhances membrane–cortex attachment primarily through increased actin polymerization at the activated site.

### E. An effective membrane chemical potential including tension and adhesion energy can account for tension gradient homeostasis

Given our experimental observations, we sought to formulate a biophysical model that explains how a persistent membrane-tension gradient can be maintained across the cell in the absence of large-scale lipid and actin fluxes, and thus without associated frictional dissipation. To this end, we use a coarse-grained description based on an effective free energy

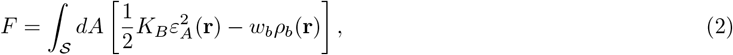

where 𝒮 is the membrane surface, *K*_*B*_ is the membrane area-stretch modulus, ℰ_*A*_(**r**) is the local elastic area strain, and 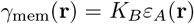 defines the corresponding in-plane membrane tension. The second term describes the energetic gain from membrane-cortex binding, with *ρ*_*b*_(**r**) denoting the local density of bound membrane–cortex binding sites and *w*_*b*_ the binding energy per binding site (*w*_*b*_ *>* 0).

To describe lipid redistribution, we derive a chemical potential associated with lipid number. In analogy to previous results (20, 21) and assuming *ρ*_*b*_ *∝ ρ*_*𝓁*_ with *ρ*_*𝓁*_ the density of lipids in the membrane, we obtain

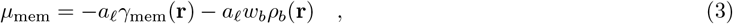

where *a*_*𝓁*_ is the resting area per lipid molecule in the membrane. Here, we retain only terms to zeroth order in the area strain. This form of the chemical potential reflects that lipid addition locally reduces membrane elastic energy, while membrane–cortex binding lowers the free energy through associated binding interactions. We note that, interestingly, this chemical potential is proportional to the apparent membrane tension, i.e. the tension that can be measured by membrane tether forces.

Motivated by the observed increase in cortical f-actin (Fig. 2b) and tether forces (Fig. 3b), we attribute a localized increase *δρ*_*b*_ in membrane–cortex attachment density to localized RhoA activation in the stimulated cell half, such that 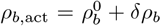. In contrast, for the non-activated half, where RhoA is not stimulated, we assume the attachment density remains at its basal value 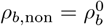. Furthermore, based on our FliptR measurements, we assume that the in-plane membrane tension in the activated half remains at its baseline value, 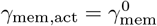.

Our FliptR lifetime measurements (Fig. 1c) suggest that after the onset of optogenetic activation, the membrane relaxes to a quasi-steady state within tens of seconds in which lateral lipid fluxes vanish and the thermodynamic driving force for lipid flow decays to zero. This corresponds to a spatially uniform membrane chemical potential of lipids, i.e. *µ*_mem_ = const. and thus *µ*_mem,act_ = *µ*_mem,non_. Correspondingly, we obtain

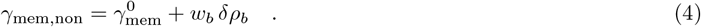

Thus, an increase in membrane–cortex attachment density on the activated side is balanced by an elevated in-plane membrane tension on the non-activated side, in agreement with our experimental observations, see Fig. 1c and 3b. Moreover, because the apparent membrane tension is proportional to the lipid chemical potential in this description, rapid equilibration of *µ*_mem_ is consistent with rapid equilibration of apparent membrane tension across the cell. In summary, the concept of a spatially uniform lipid chemical potential in steady state provides a framework for understanding how variations in membrane–cortex adhesion can maintain membrane-tension gradients across the cell, analogous to the way osmotic gradients sustain hydrostatic pressure differences in hydrodynamic systems (14, 22).

## III. DISCUSSION

Membrane tension is a major regulator of cellular signaling and of the transmission of extracellular cues (1–3). Despite its biological importance, substantial debate remains over the extent to which membrane tension gradients can be maintained within cells and over the timescales on which such differences are dissipated by tension propagation (7, 9). Previous studies have addressed this question primarily using tether-force measurements. These studies showed that direct perturbation of the plasma membrane by tether pulling induces a local increase in membrane tension, but does not lead to long-range tension propagation across the cell on timescales of hundreds of seconds (7, 9). To account for this observation, it has been proposed that frictional coupling between the membrane and the cytoskeleton strongly dampens membrane tension propagation (7). Surprisingly, however, local perturbations of the actin cortex induced by optogenetic activation of RhoA via Opto-LARG were shown to trigger rapid, cell-wide changes in tether force on timescales of only a few seconds (9). Correspondingly, de Belly et al. (9) proposed that membrane tension can propagate rapidly across the cell when forces are applied through the cortex. Together, these findings point to an apparent discrepancy between membrane tension propagation following direct membrane pulling and that following localized cytoskeletal perturbation.

To further examine this discrepancy, we measured membrane tension using an alternative approach based on the fluorescent, tension-sensitive membrane probe FliptR, whose fluorescence lifetime positively correlates with in-plane membrane tension (11, 12). Using this method, we find that localized optogenetic activation of RhoA gives rise to sustained, non-uniform FliptR lifetimes across the cell, indicating the presence of a cell-scale membrane tension gradient with elevated tension on the non-activated side. Notably, this gradient persists for hundreds of seconds, consistent with the slow tension relaxation reported by Shi et al. (7).

At first glance, these observations appear to be in disagreement with the results of de Belly et al. (9), who inferred rapid, cell-wide tension propagation from tether-force measurements. However, tether forces report an apparent membrane tension, *γ*_app_ = *γ*_mem_ + *γ*_adh_, which includes contributions from both the in-plane membrane tension *γ*_mem_ and the membrane–cortex adhesion energy *γ*_adh_. Our measurements show that tether forces increase locally upon RhoA activation, indicating an increase in *γ*_app_, while FliptR measurements reveal no corresponding increase in inplane membrane tension (see Fig. 1 and 3).Our findings therefore suggest that the increase in tether force at sites of localized RhoA activation is primarily driven by enhanced membrane–cortex adhesion, rather than by changes in in-plane membrane tension. Consequently, changes in tether forces following optogenetic RhoA activation should not be directly interpreted as changes in membrane tension itself, but rather as changes in the composite quantity *γ*_app_. Our measurements further suggest that the formation of a cell-scale membrane tension gradient upon localized RhoA activation depends on an intact actin and microtubule cytoskeleton (see Fig. 1). This process is accompanied by a cytoskeletal polarization, characterized by increased cortical f-actin at the site of optogenetic RhoA activation and an accumulation of microtubules on the opposite, non-activated side (see Fig. 2). We speculate that this cytoskeletal polarization amplifies the initial biochemical and mechanical asymmetry through the following mechanism; a well-established driver of cell polarization is the mutual antagonism between RhoA and Rac1, which promotes their spatial segregation into contractile and protrusive domains (23, 24). In migrating cells, Rac1 activity is typically enriched at the leading edge, where it drives actin polymerization and protrusion, while RhoA activity dominates at the rear, promoting contractility. This reciprocal inhibition creates a bistable system capable of amplifying small initial asymmetries into stable polarity states (23, 24). In our system, localized RhoA activation may suppress Rac1 activity at the stimulated site while indirectly promoting Rac1 activity on the opposite side, consistent with previously reported long-range partitioning of Rho-family GTPases. Although Rac1 dynamics were not directly measured here, such antagonistic coupling provides a plausible biochemical framework for reinforcing mechanical asymmetry by linking local cortical reinforcement to distal membrane and cytoskeletal remodeling. In particular, increased Rac1 activity at the non-activated side could account for the observed over-proportional increase in microtubule polymerization (25). Conversely, enhanced microtubule density is known to further promote Rac1 activation (26), giving rise to a positive feedback loop between Rac1 signaling and microtubule organization. Through Rac1-mediated inhibition of RhoA, this feedback is expected to further sharpen the spatial segregation of RhoA activity, thereby amplifying the initial asymmetry induced by optogenetic activation.

Our findings complement recent work by García-Arcos et al. (27), who demonstrated that adherent, front–back polarized cells can maintain steady-state membrane tension gradients through local force balance between actin-driven protrusion and adhesion-mediated anchoring. While their study focused on the maintenance of pre-existing asymmetries in polarized cells, our results demonstrate how such gradients can be dynamically generated from an initially isotropic state through localized signaling. Together, our results support a model in which localized RhoA activation generates spatially patterned membrane tension through cortical reinforcement and increased membrane–cortex coupling, while long-range redistribution is shaped by microtubule dynamics and intracellular RhoA–Rac1 polarity circuits. In this context, a coarse-grained membrane chemical potential of membrane lipids provides a useful alternative conceptual framework, predicting local variations in membrane tension arising from spatial differences in membrane–cortex adhesion, analogous to osmotic pressure gradients in fluids.

Collectively, our findings further shift the conceptual framework of membrane tension from one centered on rapid dissipation toward one in which cells actively generate and maintain spatially patterned mechanical states. While prior studies established that the cortex enables long-range force transmission in tether-based measurements, our work demonstrates that such transmission can give rise to stable, in-plane tension gradients rather than complete mechanical equilibration.

## IV. MATERIALS AND METHODS

### A. Cell culture

In this study we used wild-type and transgenic HeLa Kyoto cells expressing photo-activatable RhoA (opto-RhoA-mCherry) (17). The cells were cultured in Dulbecco’s Modified Eagle Medium (DMEM) supplemented with 10% fetal bovine serum (FBS), 1% penicillin/streptomycin. For transgenic HeLa cells expressing opto-RhoA, geneticin was added to the medium at a concentration of 0.4 mg/mL. Cell culture flasks were maintained at 37°C in a 5% CO_2_ environment. Cells were passaged every 2 − 3 day when they reached 60-80% confluency. The transgenic HeLa cell line was generated through lipofection using Turbofectin 8.0 (#TF81001, OriGene) according to the manufacturer’s instructions. The opto-RhoA-mCherry pcDNA3.1 vector was obtained as a gift from Brian Chow (Addgene plasmid #164472 ; http://n2t.net/addgene:164472; RRID:Addgene_164472).

### B. Measurement of FliptR lifetimes during localized RhoA photoactivation

One day before the experiments, 20,000 HeLa Kyoto cells (opto-RhoA-expressing) were seeded into an Ibidi *µ*M-Slide 8-Well glass bottom dish (1 cm^2^ growth area, Cat. No. 80827). 2 to 4 hours before measurement, the growth medium was replaced with a CO_2_-independent imaging medium, and STC (4 *µ*M) (#164739, Sigma) was added to arrest the cells in mitosis. FliptR dye was introduced at a concentration of 2 *µ*M at least 15 minutes before the start of the measurement. For control measurements, imaging protocols were applied to wild-type HeLa Kyoto cells expressing no opto-RhoA construct.

The experiments were conducted using an LSM 780 confocal microscope (Zeiss) at the CMCB light microscopy facility at TU Dresden. A Plan Apochromat 20×*/*0.8 air objective (Zeiss) was utilized with pixel number 512 × 512 and image size of 106.3 × 106.3 *µ*m^2^. During the measurements, the cells were maintained at 37°C inside an incubation chamber. For opto-RhoA-mCherry activation and FliptR FLIM imaging, a 20 MHz pulsed 473 nm excitation laser was used (imaging: 0.8%, activation: 4% laser power). Activation was performed intermittent with imaging using the ‘bleaching’ tool of the Zeiss software.

For each measurement, a region of interest with 2 *−* 5 mitotic cells with bright opto-RhoA-mCherry fluorescence was chosen and the focus was adjusted to the equatorial cross-section of the mitotic cells. Then, a short preliminary time series was recorded to verify that opto-RhoA is photo-activatable upon blue light exposure in selected cells; to this end, we imaged cells in a time series with a 561 nm laser at a time interval of 5 s. Localized photo-activation was realized by ‘bleaching’ with the 473 nm laser in a rectangular region of ≈8 ×20 *µ*m covering a part of the cell cortex that had been selected for RhoA activation for each selected mitotic cell, see Fig. 1 b. Activation was repeated after acquisition of each image in the time series. Cells were selected as suitable for actual measurement if increased mCherry-fluorescence was emerging upon activation at the cell boundary in the activation region within ≈1 min.

Then a 2nd timeseries of the same region was recorded performing the actual FLIM imaging (imaging settings: excitation with a 20 MHz pulsed 473 nm laser, bi-directional scanning, pixel size of 0.21 *µ*m, frame averaging of 2 frames, pinhole size of 2.92 Airy units, pixel dwell time of 0.64 *µ*s, a time interval of 10 s for 12 initial frames without opto-RhoA activation, subsequent images recorded at a time interval of ≈5 s with intermittent opto-RhoA activation spanning a time interval of at least 400 s). In the Becker&Hickl FLIM software, FLIM images were captured as “.sdt” images in FIFO Imaging mode. In total, 150-180 cycles were recorded with a collection time of 10 s (repetition time: 15 s, display time: 20 s, gain: 2).

Where indicated, cytoskeletal drugs were added 15-20 minutes before the imaging start. Final concentrations of cytoskeletal drugs used: PN-Blebbistatin (20 *µ*M, #DR-N-111 Optopharm), Nocodazole (10 *µ*M, #31430-18-9 Sigma), and CytochalasinD (4 *µ*M, #11330, Cayman Chemical). Stock concentration of drugs were as follows: PN-Blebbistatin: 20 mM, CytochalasinD: 4 mM, and Nocodazole: 10 mM. Correspondingly, DMSO concentrations in the medium were smaller or equal to 0.1% v/v. Each condition was measured in at least two independent replicates.

For image analysis and FliptR lifetime quantification, first the cell periphery of individual cells was identified as an annular region of high FliptR fluorescence intensity with a width of ≈ 40% of the cell radius centered at the cell periphery. Only photons from this peripheral annulus were analyzed restricting analysis to signal from the plasma membrane. The activated and non-activated half of the cell were manually selected by drawing a region of interest that captures the activated half of the cell using the ‘drawrectangle’ function of matlab. By focusing solely on the lifetimes of photons from the activated and non-activated half of the cortical region, FLIM lifetime data were analyzed for individual cells over time. Consequently, the time dynamics of fitted FliptR lifetimes and cell radii following localized RhoA photoactivation were monitored over time. Lifetimes of each cell were normalized by dividing through the median of the lifetimes measured during the first five time points (before activation started).

### C. Measuremenet of f-actin and microtubule concentration changes upon localized activation of RhoA

Seeding was done as described in Section IV B. The experiments were conducted using an LSM 780 confocal microscope (Zeiss) at the CMCB light microscopy facility at TU Dresden. For f-actin imaging, a C-Apochromat 40 ×*/*1.2 water Korr FCS M27 objective (Zeiss) was used. For microtubule imaging, a Plan Apochromat 20×*/*0.8 air objective (Zeiss) was chosen. During the measurements, the cells were maintained at 37°C inside an incubation chamber. f-actin was visualized through SiR-actin (SPY620-actin). Microtubules were visualized through Tubulin Tracker™ Deep Red fluorescence. Either dye, was added to the medium one hour before imaging start at a final concentration of 0.5 *µ*M.

For both measurements, a region with 2-3 cells was selected first and tested for photo-activatibility. Then, the timeseries imaging to observe cortical actin flow was done with the following imaging settings: bi-directional scanning, 512 *×*512 pixels^2^, pixel size of 0.1 *µ*m, frame averaging of 2 frames, pinhole size of 2.3 Airy units, pixel dwell time of 1.27 *µ*s, 240 cycles, a time interval of 10 s for 12 initial frames without opto-RhoA activation, subsequent images recorded at a time interval of ≈5 s with intermittent opto-RhoA activation.

To monitor f-actin distribution and potential actin flow along the cortex, we generated kymographs of SiR-actin fluorescence along the cortex using the KymographBuilder plugin in Fiji. To quantify f-actin and microtubule concentrations, we used Fiji to quantify SiR-actin and Tubulin Tracker^*T M*^ fluorescence intensity in selected regions of interest, see Fig. S1.

### D. AFM membrane tether pulling

Seeding was done as described in Section IV B. The next day cells were arrested in mitosis using 4 *µ*M S-trityl-L-cysteine (STC) (#164739, Sigma) and kept in incubation in a CO_2_-independent imaging buffer at 37°C for 2 hours before starting the measurement. For drug-treated conditions, drugs were added at specified concentrations to the medium approximately 15 min before the measurement.

Measurements were performed with an AFM Nanowizard 4XP (Bruker) mounted on an inverted LSM900 confocal microscope (Zeiss) at the PoL light microscopy facility, TU Dresden. The experiments used a Plan Apochromat 20×*/*0.8 air objective (Zeiss) to locate the cells with transmitted light imaging.

MLCT-C AFM cantilevers with pyramidal tip (nominal spring constant of 0.005 *−*0.02 N/m, Bruker) were employed for the membrane tether-pulling assay. Before the measurement, cantilevers were plasma cleaned for 2 minutes and then incubated in a PBS solution containing 2.5 mg/ml FITC-conjugated Concanavalin A (Sigma, #C7642) for 30 minutes at room temperature.

The cantilever’s spring constant was calibrated on each measurement day using thermal noise analysis (JPK-SPM software). During the measurement, the cells were maintained at 37°C using a Petri dish heater (JPK Instruments). Cells were measured individually; first, a mitotic cell was selected based on its spherical shape. The cantilever was then approached towards the cell, positioning the tip in the center of the cell. Then, a scan was performed in a 4 × 4 grid pattern (with an edge length of 0.5 *µ*m) at the center of the mitotic cell. For each grid point, tether-pulling commenced by lowering the cantilever onto the cell at a speed of 1 *µ*m/s until a set-point force of 1 nN was reached. The cantilever was then held at a constant height for 1 *s* before retracting by 5 *µ*m at the speed of 1 *µ*m/s for different cytoskeletal drug-treated conditions.

During cantilever retract from the cell surface, AFM forces decline and eventually become negative due to remaining adhesive contact between the cantilever and membrane tethers, see Fig. 3a. Lifting the cantilever further, membrane tethers gradually detach giving rise to sudden force jumps, see red ellipses in Fig. 3a. Each condition was measured in at least three independent replicates. The order in which conditions were measured was varied between sessions to minimize potential biases from prolonged incubation times.

Throughout the measurements, the force on the cantilever was continuously recorded at a frequency of 10 Hz. This data was subsequently used to measure tether-pulling forces using the ‘step fit’ analysis module of the JPK-SPM Data analysis software, which focused on positive force steps. In Fig. 3b, each cell gave rise to one data point in the statistics by calculating the median force step magnitude measured on this cell.

On each cell, first one sequence of tether forces was recorded before the initiation of optogenetic activation (‘bef. act.’). Then, confocal imaging of opto-RhoA-mCherry was started (561 nm laser for excitation) focusing on the topmost region of the cell. Imaging was alternated with intermittent optogenetic activation in a disk-shaped activation region of diameter ≈10 *µ*m through scanning with a 488 nm laser (‘Bleaching’ mode of the Zeiss software with bleaching after each time point). After approximately 100-200 s of sustained activation, another sequence of tether force measurements was recorded (‘aft. act.’) while activation continued.

### E. Cortical tension measurement using AFM

One day before the experiments, 10,000 HeLa Kyoto cells expressing opto-RhoA-mCherry were seeded into a silicon cultivation chamber (growth area: 0.56 cm^2^, from ibidi 12-well chamber) attached to 35 mm glass-bottom dishes (FD35-100, Fluorodish). The silicon inserts were removed 2 −8 hours before the measurements, and the growth medium was replaced with a CO_2_-independent imaging medium composed of DMEM (12800-017, Invitrogen) with 4 mM NaHCO3, buffered with 20 mM HEPES/NaOH at pH 7.2, and supplemented with 10% FBS. At this stage, S-trityl-L-cysteine (STC) (#164739, Sigma) was added to the medium at a concentration of 4 *µ*M to arrest the cells in mitosis. The cytoskeletal drugs were introduced to the cells at least 15 minutes before the measurement began. Measurements were performed with an AFM Nanowizard 4XP (Bruker) mounted on an inverted LSM900 confocal microscope (Zeiss) at the PoL light microscopy facility, TU Dresden. The experiments used a Plan Apochromat 20 ×*/*0.8 air objective (Zeiss) to locate the cells with transmitted light imaging. The cells were maintained at 37°C using a Petri dish heater (JPK Instruments). Tipless AFM cantilevers (HQ: CSC37/tipless/No Al, nominal spring constant of 0.3 −0.8 N/m, Mikromasch) were prepared for the parallel confinement assay by adding a wedge made of UV-curing adhesive (Norland Optical Adhesive 63, Norland Products) to correct for a 10° tilt. On each measurement day, the spring constant of the cantilever was calibrated using thermal noise analysis (JPK-SPM software).

For measurement of cortical tension changes due to localized opto-RhoA activation, cells were measured one by one. First, an opto-RhoA activatable mitotic cell (spherical cells) was selected. Then, an approach was made with the cantilever to the bottom of the dish, in close proximity to the chosen cell, to determine the relative z-position of the dish bottom. The cantilever was subsequently retracted by approximately 18 *µ*m and gently positioned over the mitotic cell to achieve parallel plate confinement. The step for uniaxial cell compression was initiated in the AFM software, lowering the wedged cantilever by 3 *µ*m at a speed of 0.2 *µ*m*/*s. The cantilever was maintained in the lowered position for ≈100 s before initiating confocal imaging of opto-RhoA-mCherry in the equatorial plane of the cell (561 nm laser) with intermittent activation with a 488 nm laser in a rectangular activation area, see also activation as described in IV B. Activation was maintained for at least 200 s until the measured AFM force reached a new equilibrium value. Then, optogenetic activation was stopped and force equilibration was recorded for another *≈* 100 s. Then, the cantilever was retracted and lifted and the measurement was stopped.

During the measurements, the force on the cantilever and its height relative to the dish bottom were continuously recorded using software provided by the AFM manufacturer (JPK Instruments). These data were then used alongside the confocal images to determine the cell’s surface area, volume, and cortical tension as previously described (16). Then, the intracellular hydrostatic pressure Δ*p* was inferred from cortical tension using the Young-Laplace equation which we approximated as Δ*p* = *γ*_cortex_(1*/R*_cell_ + 2*/h*) for confined cells, where *h* was the confinement height and *R*_cell_ the measured cell radius during confinement (14). Pressure values were determined for 3 time points, before, during and after optogenetic activation using the corresponding value of equilibrated AFM forces.

## ACKNOWLEDGMENTS

We thank Michael Schlierf and James Saenz for the good discussions on the project. Furthermore, we thank the Deutsche Forschungsgemeinschaft (DFG, German Research Foundation) for financial support through the Heisenberg program, project number 495224622/FI 2260/8-1 (EFF) and the grant FI 2260/5-2 (EFF). In addition, the authors thank the CMCB Light Microscopy Facility, the PoL Light Microscopy Facility for their excellent support.

## DATA AVAILABILITY STATEMENT

All data and code generated during this study are available from the corresponding author upon reasonable request.

## AUTHOR CONTRIBUTIONS

Y.A.P. performed the experiments; Y.A.P. and E.F.-F. jointly designed the experiments and analyzed the data. Y.A.P. and E.F.-F. wrote the manuscript.

## Supplementary Material

**Figure S1.**
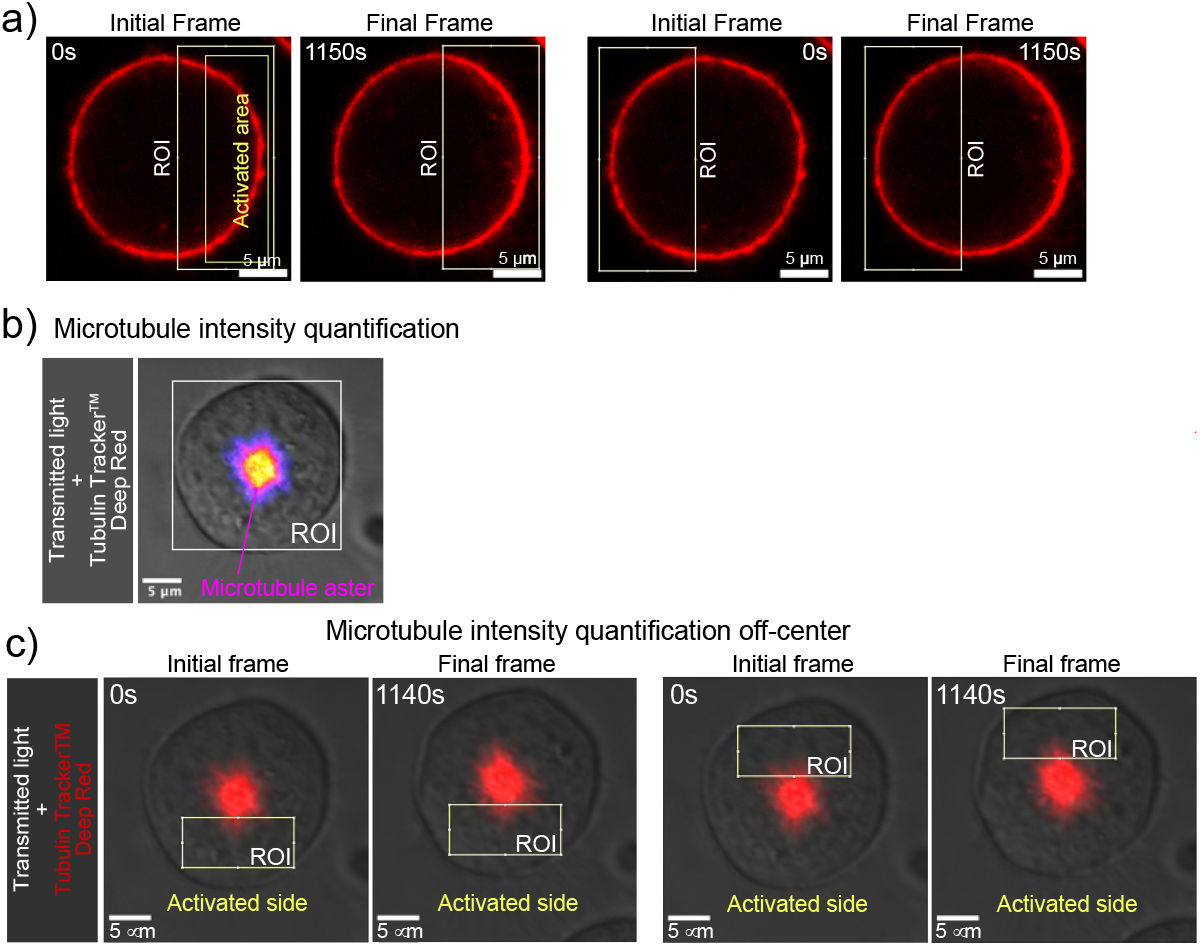
Image-analysis workflow for quantifying actin and microtubule polymerization. a) Quantification scheme for cortical f-actin changes, see Fig. 2a,b. b) Quantification scheme for total microtubule intensity in the cell, see Fig. 2g,h. c) Quantification scheme for of off-center microtubules adjacent to the central aster both on the activated and non-activated sides, see Fig. 2h..

